# SNP2SIM: A modular workflow for standardizing molecular simulation and functional analysis of protein variants

**DOI:** 10.1101/457192

**Authors:** Matthew D. McCoy, Vikram Shivakumar, Sridhar Nimmagadda, Mohsin Saleet Jafri, Subha Madhavan

## Abstract

Molecular simulations are used to provide insight into protein structure and function, and have the potential to provide important context when predicting the impact of sequence variation on protein function. In addition to understanding molecular mechanisms and interactions on the atomic scale, translational applications of those approaches include drug screening, development of novel molecular therapies, and treatment planning when selecting targeted therapies. Supporting the continued development of these applications, we have developed the SNP2SIM workflow generates reproducible molecular dynamics and molecular docking simulations for downstream functional variant analysis. Three modules execute molecular dynamics simulations of solvated protein variant structures, analyze the resulting trajectories for unique structural conformations, and bind small molecule ligands to representative variant scaffolds. In addition to availability as a command line workflow, SNP2SIM modules are also available as individual apps on the Seven Bridges Cancer Genomics Cloud.

## Background

Molecular simulation is a powerful tool used by computational biologists to analyze the relationship between protein structure and its functional properties. Ranging from high throughput drug screening to focused characterization of protein conformational dynamics, the creative analysis has several translational applications. Large libraries of drug candidates can be evaluated to produce novel targeted therapeutics, and insight into specific molecular interactions between effective drugs and their protein targets aids the design novel molecules [1, 2]. An advantage of the computational simulations is the ability to probe how variation in the protein sequence alters those molecular interactions, and can be extended to the development of therapies targeted at specific sequence variants [3-6]. In addition to drug discovery and design, the insight can be further extended to inform treatment planning when selecting an optimal targeted therapeutic strategy [7].

Due to an inherent tradeoff between resolution and computational requirements, molecular simulations can be divided between approaches which only simulate a fraction of the overall molecule and those which explicitly consider all atomic interactions occurring within the molecule. Coarse grained methods which do not explicitly consider the internal interactions occurring within the protein backbone used to address the enormous search space that must be interrogated when predicting how two molecules interact [8]. For example, predicting how well a small molecule ligand will bind to a target protein depends on the sum total of all the individual atomic interactions. Depending on the chemical nature of the ligand, the conformational diversity can be quite large due to rotation around individual bonds and limited steric constraints of a single ligand molecule. Furthermore, the protein surface represents a large area of potential interactions and exponentially increases the degrees of freedom which must be explored when identifying an optimally bound structure. In order to simplify the search for optimized protein:ligand conformations and to simulate high throughput binding of large libraries of low molecular weight ligands, coarse grained docking methods will typically only model the flexibility of the ligand and a small number of interacting protein residues within a defined area of a rigid protein structure [8].

While the liberties taken by these types of simulations allow for a greater throughput, they fail to account for internal protein dynamics which may play a significant role in the interacting complex. All-atom molecular dynamics (MD) simulations explicitly account for atomic interactions occurring within a molecular system and provide a way to understand the overall conformational flexibility and structural dynamics [9]. However, even systems consisting of a small, solvated protein contain tens to hundreds of thousands of atoms and each simulation step requires a summation of all the forces acting on each. Even on high performance computational infrastructures, simulation runs can easily last weeks to generate usable results. The increased computing cost is offset by its unique insight and characterization of functionally relevant protein dynamics.

Both approaches find utility in specific applications, and their individual strengths are leveraged to understand the impact on protein sequence variation on small molecule binding. When a new amino acid is specified by a change to the genomic sequence, the change in the residue side chain has the potential to alter the functional interactions with a small molecule. If the change occurs within the defined search space of a coarse grained binding simulation, the new interactions can be simulated directly. Typically, the structures used for binding simulations are derived from x-ray crystallography, but simply swapping out amino acid side chains in the intersecting residues may not fully account for the structural differences of the protein variant. Since the protein backbone is treated as a rigid scaffold, the resulting predicted binding characteristics do not account for those subtle changes in the backbone geometry and could have a large influence on the results. Furthermore, these methods have nothing to offer if the variation occurs outside of the defined search space, especially those amino acids which are buried within the folded protein structure. MD simulations can address this limitation by comprehensively sampling the conformational landscape of a protein variant to generate characteristic scaffolds for downstream small molecule docking.

Since a protein variant can alter the functional interaction with therapeutic molecules, predicting how small molecules will bind to protein variants has a significant application in personalized medicine. Not only can simulation results be used in the development of targeted therapies, it could also be informative in the selection of second line of therapy once drug resistance has emerged. As the application of molecular profiling and sequence analysis continues to gain a foothold in clinical decision making, a well-defined, user friendly simulation workflow and methodology would be an important tool for translational computational biology. To that end, we present SNP2SIM (**Figure 1**), a scalable workflow for simulating the impact of protein sequence variation on binding to small molecule ligands.

**Figure 1.**
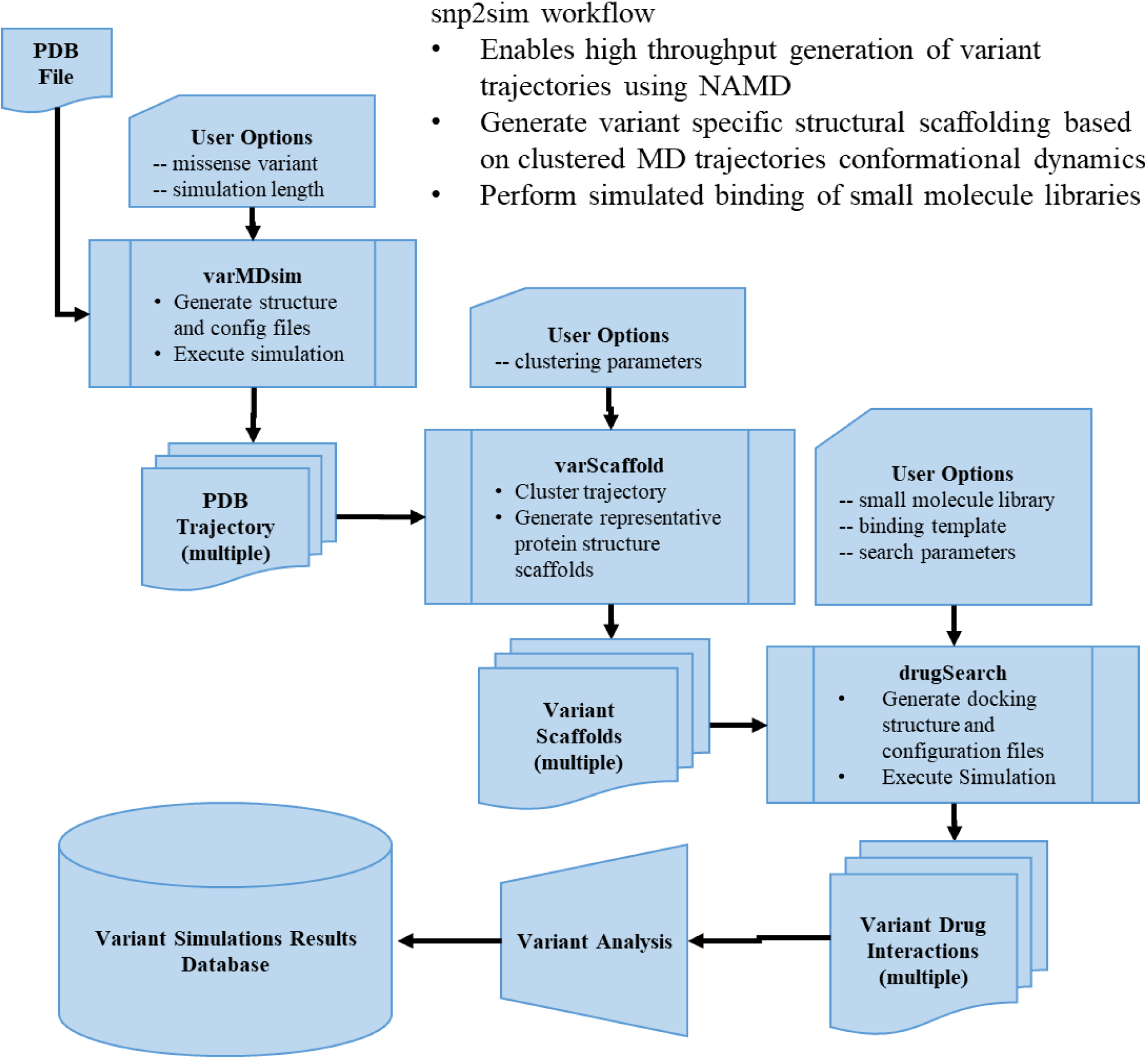
The SNP2SIM workflow contains 3 functional modes; varMDsim generates molecular dynamics trajectories using NAMD, varScaffold clusters the resulting trajectories into variant specific representations of the structural variation, and drugSearch binds a library of low molecular weight ligands to each variant scaffold.

## Implementation

At its core, SNP2SIM is a modular set of simulation and analysis tools wrapped in a command line python script. The three functional modules correspond to the molecular dynamics simulation of a protein variant, conformational analysis of molecular dynamics trajectories, and small molecule docking to variant specific structural scaffolds. The workflow controls the generation of tool specific preprocessing and analysis scripts, configuration files, and file structure based on an initial PDB formatted protein structure file, and executes the simulation software. The command line implementation of SNP2SIM is available for download (https://github.com/mccoymd/SNP2SIM), and the varMDsim, varScaffold, and drugSearch modules are also available as apps on the Seven Bridges Cancer Genomics Cloud [10] (http://www.cancergenomicscloud.org/).

### varMDsim

With the minimal input of a PDB formatted protein structure file and simulation time in nanoseconds, the varMDsim module will generate a solvated, ionized water box, generate the configuration files for the all-atom simulation, and compile the results for downstream analysis. Specifying an amino acid variant will automatically mutate the input structure prior to solvation. The varMDsim module utilizes versions of Visual Molecular Dynamics (VMD) [11] and Nanoscale Molecular Dynamics (NAMD) [12] installed on the user’s system, and the CHARMM36 topology and parameters [13] and simulation configurations files are hardcoded into the workflow, standardizing the resulting simulation for reuse and promoting the reproducibility of the computational simulations.

### varScaffold

The simulation trajectories are analyzed using the varScaffold module to produce characteristic structures of variant proteins. The user supplied clustering parameters specify how the protein structures are first aligned, and then clustered based on root-mean-square deviation (RMSD), using VMD’s Atom Selection Language and clustering plugin. This separate alignment and clustering parameters allow for the investigation into protein specific features of interest. For example aligning an entire protein structure by its backbone residues and clustering by the geometry of the binding pocket captures specific structural variation impacting the functional interaction with a ligand. Similarly, this can be used to measure internal dynamic behavior, such as the motion of a disordered region or positions of internal structural features. Representative PDB structures are generated for each unique cluster that is populated by at least 10% of the simulated trajectory at a given RMSD cluster threshold. The varScaffold module will accept multiple PDB or DCD formatted trajectory files generated through parallel execution of the varMDsim module.

### drugSearch

The drugSearch module uses AutoDock Vina [14] to bind a predefined library of low molecular weight molecules into the variant scaffolds. This requires the user to supply a PDB formatted protein structure, and an associated parameter file that defines the search space for ligand binding. Additionally, the user can specify a set of residues within that search space model with flexible sidechains. Variant scaffolds are aligned to the reference coordinates, and the associated configuration files are generated for each ligand in the drug library. General analysis tools included along with the SNP2SIM package include bash scripts to compile the quantified AutoDock Vina results from multiple files, generate PDB formatted files of the ligand and flexible side chain orientations, and to visualize the relative binding affinity between wildtype and variant structures.

### Case Study: PD-L1 small molecule inhibitors

The immunomodulatory protein PD-L1 was used to demonstrate the application of the SNP2SIM workflow to drug development in personalized medicine. Development of small molecule inhibitors has clinical applications, and a number of molecules are currently being investigated for therapeutic use in cancer. To understand how these molecules may differentially bind to variants of PD-L1, known mutations in the binding domain were processed through the SNP2SIM workflow. The initial starting structure used the Ig-like V-type domain from PDB: 4Z18, and 500 ns of simulation were generated for each protein variant. Variants were selected based on their occurrence in PD-L1 expressing cell lines as well as those most commonly occurring across all cancer types (L53P, V68L, R86W, L94M, G95R, A97V, M115T). Variant trajectories were aligned using the entire domain backbone and clusters were defined using a 0.7 RMSD cluster threshold for the backbone atoms in residues interacting with low molecular weight inhibitors in PDB crystal structures(cite) (Residues 19, 20 54, 56, 66, 68, 115, 116, 117, 121, 122, 123, 124, 125). These same interacting residues were also modeled with flexible side changes when bound to a library of 17 small molecule ligands. The SNP2SIM workflow was run using the Seven Bridges Cancer Genomics Cloud infrastructure (cite).

## Results

The SNP2SIM workflow enables the efficient parallelization of the computationally intensive molecular dynamics simulations. Variant structures of PD-L1 were simulated for 5 independent runs of 100 ns each (total 50 ns), and the resulting trajectories were combined for downstream analysis. The RMSD of both the domain backbone and small molecule binding residues (**Figure 2**), show the variants all maintain a relatively stable conformational population. Despite the overall lack of molecular motion on a global scale, the results show the variant structures behave differently relative to wildtype. This is reflected in the deviation of the entire PD-L1 domain backbone, which is even more pronounced when only considering the residues which interact with small molecule inhibitors (**Figure 3**).

**Figure 2.**
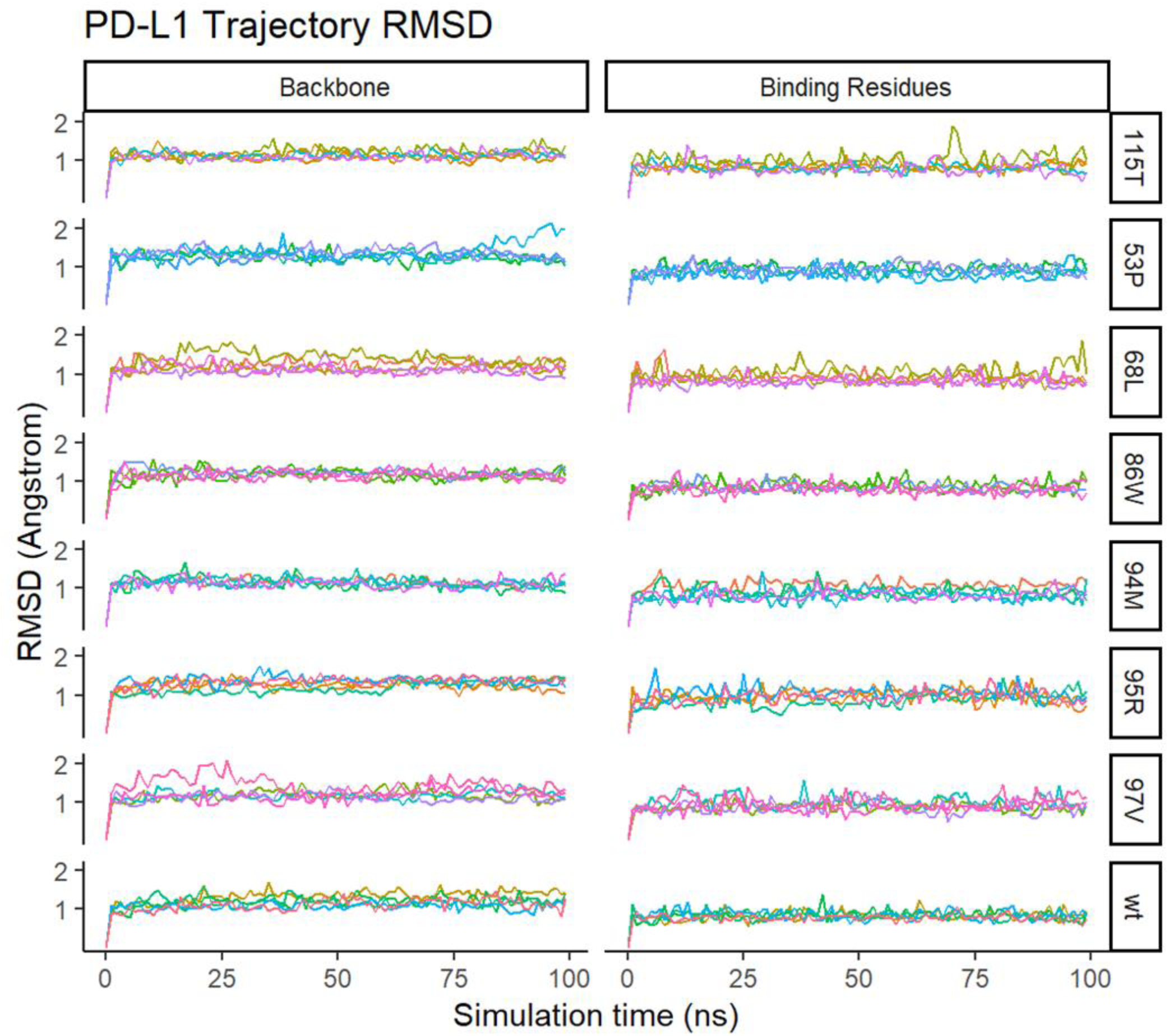
The SNP2SIM results from the varMDsim module. Each color represents an independent 100 ns NAMD simulation of the solvated PD-L1 variant structure (5 per structure variant). Root-mean-squared deviation (RMSD) of the domain backbone (alignment residues) and binding show (clustering residues) reveal differences in wildtype and variant conformational dynamics.

**Figure 3.**
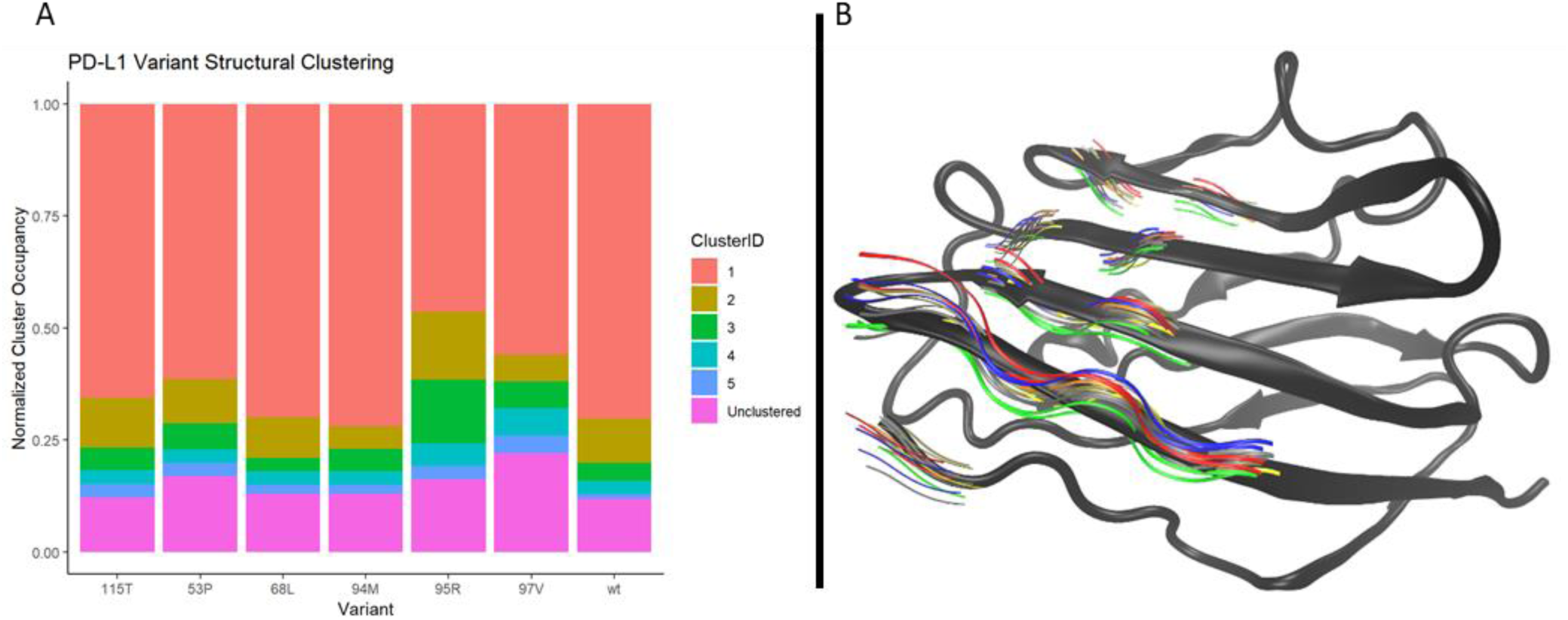
The (A) breakdown of the results from the varScaffold module of the SNP2SIM workflow show the characteristic variation induced by each missense mutation in PD-L1. Depending on the variant, molecular dynamics simulations revealed novel structural conformations (Occupancy > 10%). (B) The backbone atoms from PD-L1 binding residues from trajectory based scaffolds, where the colors correspond to the different populated clusters of a given variant relative to the crystal structure (grey).

From the initial analysis of the variant trajectories, it’s clear that certain variants induce more conformational flexibility than others. This is highlighted in the breakdown of the trajectory clustering results (**Figure 3**), where clusters were defined by the RMSD of residues involved in binding small molecules. The wildtype PD-L1 structure had two populated clusters, one just meeting the threshold of 10% of all sampled structures. Depending on the variant, the occupancy of additional clusters decreased to one (86W, 94M, and 97V), increased to three (95R), or stayed the same (53P, 68L, and 115T), illustrating the differential impact of sequence variation on the overall conformational flexibility.

The differences in flexibility translate to changes in the predicted binding affinity, and the difference can be used to predict if a given variant will be more or less likely to bind a particular ligand (**Figure 4**). Since there were two wildtype scaffolds, each was compared separately to each variant scaffold. For the same variant, the relative binding affinities are largely similar in direction and magnitude for both the wildtype conformations. But it’s not always the case, and close inspection of instances where the pattern diverges had the potential to yield significant insight into the functional nature of the protein. The same applies to differences between input scaffolds for individual variants, where the inhibitory function of certain small molecules may be related to differential binding to conformational populations.

**Figure 4.**
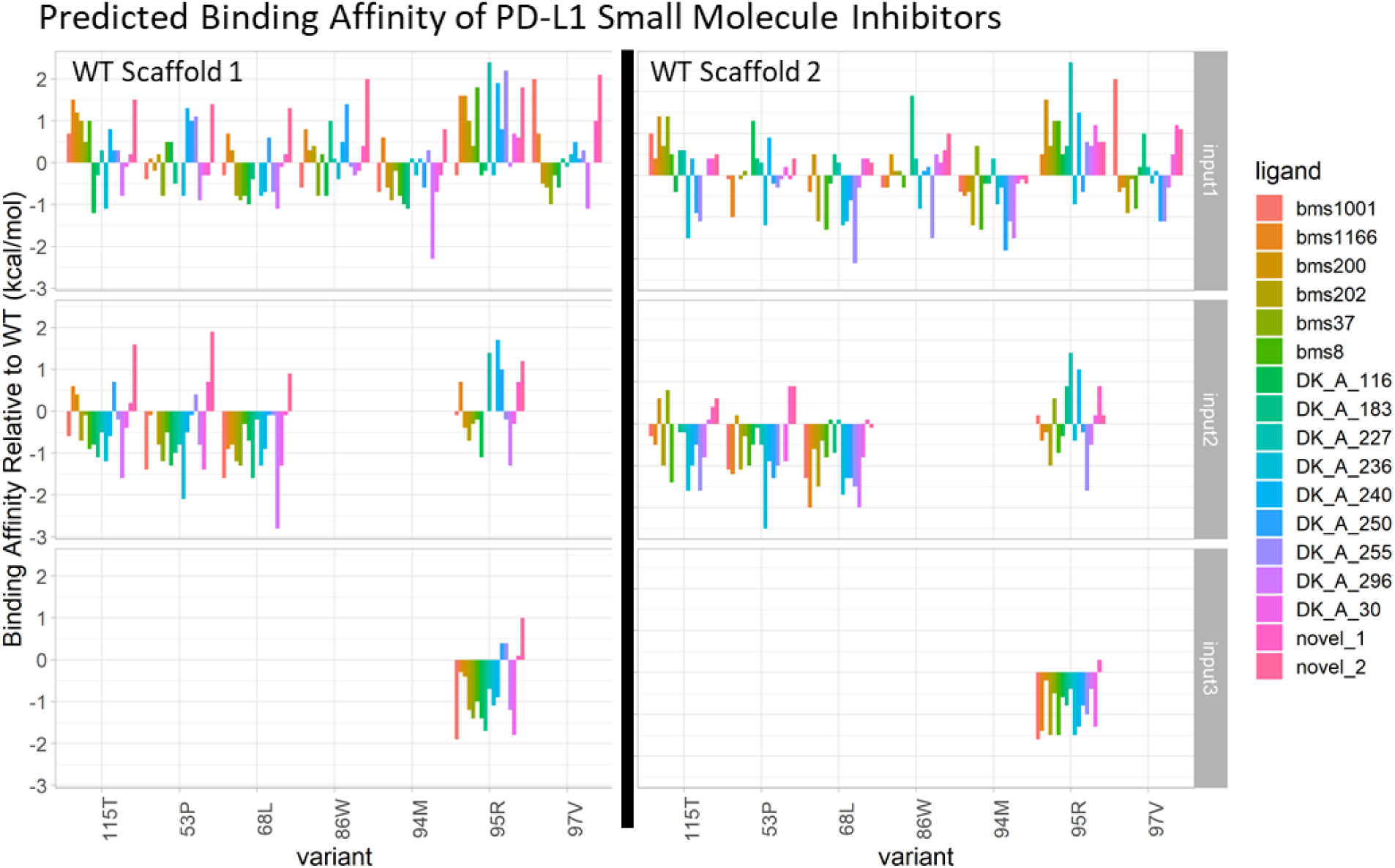
The SNP2SIM drugBinding results for trajectory-derived PD-L1 scaffolds bound to small molecule inhibitors are used to calculate the binding affinity relative to that predicted for the wildtype structure. Positive values correspond to a decreased affinity of the small molecule for the variant structure compared to the wildtype.

## Discussion

The growing prevalence of genomic testing is revealing an enormous amount of rare variants with unknown functional significance [15], underscoring the need for predictive computational analysis to determine their functional significance. This is especially true for variants which occur in proteins where the effectiveness of targeted therapeutic strategies may be disrupted. For example, missense mutations emerge in response to evolutionary pressures in a growing tumor which disrupt binding of targeted inhibitor molecules [16]. SNP2SIM enables the profiling of multiple approved inhibitors to inform the selection or design of an optimal therapy which maintains a positive clinical response [7].

By simulating the variant specific contributions to the overall protein conformational dynamics and ligand binding, the unique impact of a variant can be quantified even when the mutated residues do not occur at the interaction interface. As seen in **Figure 3**, the proportions populations of distinct protein structures is impacted in a variant specific manner. Even for the wildtype structure, two populated conformations were identified which show slightly modified geometries of the protein backbone found in the crystal structure. The results of small molecule docking show the different scaffolds bind to the small molecule ligands with different affinities (**Figure 4**). This additional information will ultimately produce more robust analysis and improve predictive models used for downstream drug development, design, and utilization.

The widespread use of molecular simulations to generate predictive data, and the insight it can provide to understanding the functional changes of protein sequence variants, is rate-limited by computational costs and scale of potential variation. Both of these barriers are being overcome through access to cheap cloud computing and the development of reproducible workflows. And while a lot has been done to lower the barrier for novice users to access these tools through widely available infrastructure such as the NCI cloud pilots, creating an easy-to-use simulation and analysis workflow opens the doors to many researchers who would otherwise not have access. As demonstrated through the case study of PD-L1, SNP2SIM can address all these issues. The modular nature and implementation as independent apps on Seven Bridges Cancer Genomics Cloud allow for parallelization, access to high performance computing resources, and a user-friendly interface.

## Conclusions

Overall, the SNP2SIM workflow represents a higher resolution approach to the *in silico* functional predictions compared to methods that provide a limited characterization of variant pathenogencity. Not only does a simulation based approach provide detail about disruption of specific functional interactions, it can evaluate the differential impact of somatic variation on targeted therapies.

